# Magnetoencephalographic Signatures of Hierarchical Rule Learning in Newborns

**DOI:** 10.1101/2020.04.28.065482

**Authors:** Julia Moser, Franziska Schleger, Magdalene Weiss, Katrin Sippel, Ghislaine Dehaene-Lambertz, Hubert Preissl

**Affiliations:** IDM/fMEG Center of the Helmholtz Center Munich at the University of Tübingen, University of Tübingen, German Center for Diabetes Research (DZD e.V.), Tübingen, Germany, Tübingen, Germany; Graduate Training Centre of Neuroscience, International Max Planck Research School, University of Tubingen, Tubingen, Germany; Department of Obstetrics and Gynecology, University Hospital, University of Tübingen, Tübingen, Germany; Cognitive Neuroimaging Unit, CNRS, INSERM, CEA, Université Paris-Saclay, NeuroSpin center, 91191 Gif/Yvette, France; Department of Internal Medicine IV, University Hospital of Tübingen, 72076, Tübingen, Germany; Department of Pharmacy and Biochemistry, Interfaculty Centre for Pharmacogenomics and Pharma Research, University of Tubingen, Tubingen, Germany

**Keywords:** Cognitive development, magnetoencephalography, rule learning, newborns, auditory mismatch response

## Abstract

Estimating the extent to which newborn humans process input from their environment, especially regarding the depth of processing, is a challenging question. To approach this problem, we measured brain responses in 20 newborns with magnetoencephalography (MEG) in a “local-global” auditory oddball paradigm in which two-levels of hierarchical regularities are presented. Results suggest that infants in the first weeks of life are able to learn hierarchical rules, yet a certain level of vigilance seems to be necessary. Newborns detected violations of the first-order regularity and displayed a mismatch response between 200-400ms. Violations of the second-order regularity only evoked a late response in newborns in an active state, which was expressed by a high heart rate variability. These findings are in line with those obtained in human adults and older infants suggesting a continuity in the functional architecture from term-birth on, despite the important immaturity of the human brain at this age.

## 1. Introduction

Numerous experiments have illustrated the abilities of newborn human infants to recognize a variety of stimuli, some being ecologically important such as the mother’s voice (DeCasper and Fifer, 1980) or face (Pascalis et al., 1995) and smell (Marlier et al., 1998). Others being more abstract such as the native language (Mehler et al., 1988), biological motion (Simion et al., 2008) or numerical representations (Izard et al., 2009). All these abilities have been tested through the comparison of two stimuli and the elicitation of a surprise, or familiarity effect, and can be explained by local computations in specialized cortical areas leading to automatic orientation responses. Independent of the complexity of the tested stimuli, these comparisons can not inform us about the depth of processing a newborn is capable of. This aspect of the newborn’s abilities is what we question here. By means of an auditory oddball paradigm with hierarchical rules, we can investigate, whether they can not only detect first-order but also second-order regularities (see Figure 1 for the experimental paradigm). To exemplify the paradigm, we can imagine the presentation of sequences of four tones in which the first three are repeated (standard) and the last one is different (deviant; e.g. sssd sssd sssd sssd etc…). “d” induces a local error signal at the level of the sequence because it is unexpected given the previous tones (first-order regularity). However, at a more global level, “d” should be expected, given the previous sequences. At this global level (second-order regularity), “sssd” is the rule and thus the appearance of the sequence “ssss” should be surprising.

**Figure 1:**
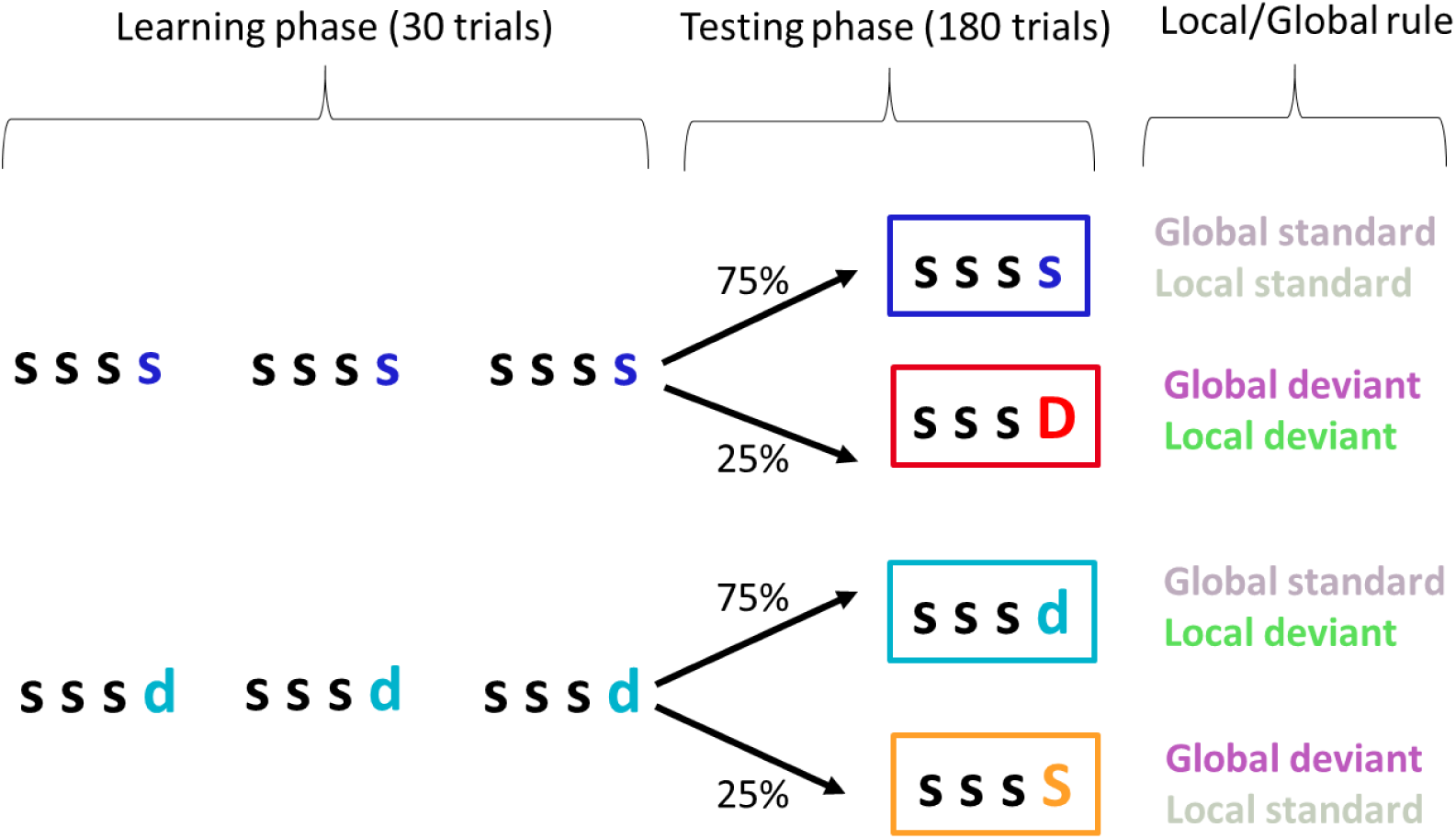
Experimental Paradigm. Each row represents one experimental block. Colors depict individual stimulus conditions. Right column describes the sequence’s role in the paradigm.

This so called “local-global” paradigm was introduced by Bekinschtein et al. (2009) who linked the ability to learn second-order regularities to conscious processing as in adult studies, unlike learning first-order regularities, learning second-order regularities could neither be observed in comatose patients (Bekinschtein et al., 2009; King et al., 2013) nor in sleeping participants (Strauss et al., 2015). Thus, learning regularities on a global level is not automatic, even in adults. Contrary to the detection of local changes based on the event frequency (or the event transition), the discovery of second-order regularities needs the formation of memory traces for chunks of elements (sequences of tones) over several seconds with an ordered comparison of each element of the actual sequence with the template in memory. The formation of memory traces plays an important role in primary consciousness which Edelman (2003) defined as the ability to learn from the environment and dynamically adapt to it.

Basirat et al. (2014) used an adapted version of this paradigm to study three months old infants. In their study, they combined auditory stimuli with matching videos to keep participants engaged and attentive. They detected signs for learning of both hierarchical rules and concluded that three months old infants consciously process those stimuli. The present study aims to unravel these processes at an even earlier age and therefore uses magnetoencephalographic (MEG) recordings with a version of the “local-global” paradigm in a sample of newborns in the first weeks of life.

We know that brain responses towards local changes can be observed even in very immature brains such as in preterms (Mahmoudzadeh et al., 2017) and fetuses (Draganova et al., 2005; Draganova et al., 2007). In infants, the so called Mismatch Response (MMR) occurs usually between 200-400ms after stimulus onset and is comparable to the adult Mismatch Negativity (Dehaene-Lambertz and Dehaene, 1994; Háden et al., 2016). In electroencephalographic recordings, this MMR is followed by a Late Slow Wave (LSW) in a time window that can vary between 680 and 1200ms (Dehaene-Lambertz and Dehaene, 1994; Friederici et al., 2002; Basirat et al., 2014). This LSW shares many common properties with the adults’ P300 and has been described as a sign of active orientation toward a new stimulus (Nelson and deRegnier, 1992) and a sign of perceptual consciousness (Friederici et al., 2002; Kouider et al., 2013). As the P300, the LSW is related to depth of processing and results by Basirat et al. (2014) showed the detection of second-order/global rule violations by the presence of a LSW.

In the context of the predictive coding framework the MMR and LSW reflect a prediction error (Friston, 2005) which refers to the discrepancy between the prediction and the actual perception. This prediction error is modulated by the formation of a memory trace (Wacongne et al., 2012) as well as by the probability of occurrence of each tone (Friston, 2005). The generation of a MMR can be described by a model including two cortical columns in the auditory cortex (Wacogne et al. 2012). The different layers of the columns represent the prediction error, the prediction and the memory trace. Their interplay lead to a specific response based on the sensory input through the thalamus. The neuronal responses we usually measure are produced in the prediction error layer. This probabilistic updating influences processing from early on and therefore, even if the MMR in the local-global paradigm is present irrespective of the global rule, its amplitude is modulated by the global rule (King et al., 2013; Basirat et al., 2014).

The LSW was so far only detected in awake and not in asleep infants (Friederici et al., 2002) but the relation between sleep, attention, novelty and late responses has not been systematically studied in infants this young. Nevertheless, it is difficult to keep infants awake during an experimental recording and to monitor their attention. In newborns periods of sleep and wakefulness are equally distributed over the day and sleep organization is very different from later on. There are only two main stages, active sleep/rapid eye movement (REM) sleep and quiet sleep whereas they spend about 50% of their sleep time in active sleep (Jenni and Carskadon, 2005). Infants fall rapidly asleep and the cycle often starts with a period of active sleep with many micro-arousals (Scher, 2008). To estimate the infant’s behavioral state, we used the heart rate variability (HRV) of our participants. Previous studies showed that infants from one week to six month of age show a higher HRV in awake state and active sleep/REM sleep compared to quiet sleep (Harper et al., 1976; Harper et al., 1982). Thereby the standard deviation of normal to normal R-R intervals (SDNN) is a valid marker to differentiate between active and quiet sleep in the first weeks (Doyle et al., 2009; Lucchini et al., 2017). This was also reported for fetal behavioral states, where HRV measured by SDNN could be used to differentiate between active and quiet states (Brändle et al., 2015).

As our paradigm and age group are not fully comparable to previous studies, we split our analysis into two parts. First, a literature driven part, where we focused on the time windows reported for the MMR (200-400ms; Dehaene-Lambertz and Dehaene, 1994; Háden et al., 2016) and the LSW, using a time window where most previous studies overlapped (800-1000ms; Dehaene-Lambertz and Dehaene, 1994; Friederici et al., 2002; Basirat et al., 2014). Thereby, we have the hypotheses to find a strong MMR towards the local rule violations (first-order regularity), which is modulated by the global rule (second-order regularity). Furthermore, a LSW like response towards all rule violations, mainly modulated by the global rule. As a second step, we added some exploratory analysis, notably to take into account the infant’s behavioral state. Here we investigated the relationship between behavioral state, quantified by HRV values, and brain responses in the local and global computations.

## 2. Methods

### 2.1. Participants

We included 33 healthy newborns and were able to record at least one complete experimental block in 27 of them (13 females, 14 males, APGAR at 10 min >8). However, after data pre- processing, only 20 participants remained with at least one experimental block of sufficient quality. Only these 20 newborns (12 females, 8 males) were included in the study (13-59 days, M=34, SD=14). The study was approved by the local ethics committee of the Medical Faculty of the University of Tübingen and the consent to participation was signed by both parents. They received 10 Euro per hour for their participation.

Prior to this study, all participants were enrolled in a fetal study using an identical paradigm to assess fetal brain responses with fetal MEG (data not shown). All 27 participants were exposed to the stimulation between one and four times (M=2.26, SD=1.35) during pregnancy.

### 2.2. Material and Design

Stimuli were two pure tones (500Hz and 750Hz, duration = 200ms), presented at 65dB sound pressure level. Tones were presented in sequences of four, separated by a 400ms inter-tone-interval (total duration of the sequence = 2 sec), each sequence was separated from the next by a 1700ms inter-stimulus-interval. Standard trials (ssss) consisted of the repetition of the same tone (i.e. the standard tone) whereas the other tone was introduced at the last position for deviant trials (sssd).

Each block started with a learning phase during which a specific sequence (ssss or sssd) was repeated 30 times. This sequence was also the frequent type of trials (75%) in the subsequent testing phase whereas the other type of sequences was randomly presented in 25% of trials. Thus two types of blocks were possible: either 75% of the sequences were standard sequences (e.g. ssss ssss ssss) with 25% deviant sequences (sssD); or 75% of the sequences were deviant sequences (e.g. sssd sssd sssd) with 25% standard sequences (sssS). Therefore, two levels of mismatches were assessed in this paradigm, either at the sequence (i.e. local) level (ssss vs. sssd) or at the block (i.e. global) level (global standard, respecting the block rule (frequent ssss or ssssd) vs. global deviant (rare sssS or sssD). Global rule violations are depicted by the upper case letter S or D (Figure 1).

Each participant was recorded with the two types of blocks, the order of blocks was counterbalanced across participants. At the beginning of each block, the learning phase implemented the global rule, followed by a testing phase with 180 trials comprising 25% global deviant trials (Figure 1). The trials in the testing phase were pseudorandomized with a minimum number of two standard trials presented between deviant trials. This resulted in an overall duration of about 13 minutes per block.

Both tones (500Hz and 750Hz) were used as standard tone or deviant tone across participants to control for a possible stimulus effect. Within one participant, standard and deviant tones were fixed.

### 2.3. Experimental procedure and recording

#### Magnetoencephalography

Magnetoencephalographic data were recorded with a MEG device which enables non-invasive measurement of heart and brain activity in both fetuses during pregnancy and infants shortly after birth (Preissl et al., 2004). For the recording of infant data, the SARA (SQUID Array for Reproductive Assessment, VSM MedTech Ltd., Port Coquitlam, Canada) system installed at the fMEG Center at the University of Tübingen was used. To attenuate magnetic activity from the environment, the device is installed in a magnetically shielded room (Vakuumschmelze, Hanau, Germany) which includes an intercom system to observe and communicate with the participant/caregiver. The system includes 156 primary magnetic sensors and 29 reference sensors. The magnetic sensors are distributed over a concave array. For infant recordings a cradle is attached to the SARA device and a parent monitors the infant inside the measurement room. For the presentation of auditory stimuli, a small child-appropriate earphone (Ear Muffins from Natus, Biologic, San Carlos, USA), was placed on one ear. In this study, all participants were placed on their right side with the earphone over the left ear. Due to the sensor design and distribution of the SARA, the head was covered by roughly 10 sensors.

#### Procedure

A time-slot of two hours was allocated to ensure full comfort of our participants. Newborns were dressed in metal-free cloths and placed in the MEG cradle on the measurement device. When the newborn was comfortable and calm, the earphone was attached and the caregiver inside the room was instructed to intervene only when absolutely necessary. The magnetically shielded room was closed and presentation of the stimuli started. Participants were monitored from outside with a camera and an intercom system. If the infant stayed calm, both recording blocks were subsequently presented, followed by a 15min silent measurement of spontaneous activity. If the infant became agitated and started crying, the measurement was stopped. If it was interrupted at the beginning of the second block, we asked the parents to calm their child and tried to continue the measurement with the second block after a short break.

### 2.4. Data Analysis

The first step in MEG data processing is the attenuation of interfering signals, especially heart signal, which is important in newborns due to the closeness of the heart relative to the MEG sensors. The magnetocardiogram was detected by template matching or using the Hilbert transform algorithm. As usually done for fetal brain analysis (e.g. Linder et al., 2014) we selected the one method which detected the heartbeats more accurately for each infant. Subsequently it was subtracted from the relevant signal, through signal space projection (Vrba et al., 2004; McCubbin et al., 2006; Wilson et al., 2008). In this step, the data were also visually checked to detect large artifacts in some channels which were removed if they were outside the area of the infant’s head. All processing steps were implemented in Matlab 16b (The MathWorks, Natick, MA).

Data were bandpass filtered from 1-15Hz and high amplitude artifacts were attenuated with an artifact block algorithm (Mourad et al., 2007). The threshold for artifact attenuation was determined by the median ±4 standard deviations of the data over all channels. If this value was higher than 2pT, the dataset was excluded due to a too-high level of noise. Threshold values for the remaining datasets varied between 0.61pT and 1.81pT (M=1.0pT, SD=0.3pT).

The continuous recordings were then segmented into epochs starting 200ms before the onset of the first tone of a sequence until 3000ms after. To keep the same number of epochs in each condition, we analyzed only the standard sequences within a block that immediately preceded a deviant one (i.e. 45 trials in each condition and in each block). Trials of each individual condition were averaged in each participant, resulting in two averages per stimulation block (ssss and sssD in the block with the ssss rule and sssd and sssS in the block with the sssd rule, Figure 1).

To optimally detect the channels containing the infant’s brain activity, a Principal Component Analysis was calculated for each average. The component coefficients were calculated to determine, which channels were the most influential for each component in the signal. The first three principal components were taken into consideration. The factor loadings were sorted to extract the five most influential channels. Those five channels were located and their location checked for plausibility. A plausible location had to fulfill three criteria: 1. Channels had a mean distance less than 10cm from each other, 2. channel locations were in the area where the newborn’s head was placed and 3. both trial types of one dataset needed a plausible location. If more than one valid location was determined, the location of the principal component with the highest explained variance was used. The method was adapted for infant recordings from Moser et al. (2019).

After data preprocessing of the 50 datasets recorded in 27 participants who had at least one complete recording block, one dataset was excluded because of no valid location and 14 datasets were marked as noisy. This led to 20 remaining participants: 15 had usable datasets for both blocks, 2 only for the block with the ssss rule and 3 only for the block with the sssd rule. 13 of the 20 participants had the 500Hz tone as standard tone and 7 the 750Hz tone (the other tone was the deviant).

#### Analysis of Heart Rate Variability (HRV)

HRV was calculated from the R peaks, previously detected during the data cleaning of the magnetoencephalogram. As HRV value, we computed the standard deviation of normal to normal R-R intervals (SDNN) in each recording block and each newborn, using in house algorithms, described in Mat Husin et al. (2020).

### 2.5. Statistical analyses

In order to test the processing hierarchy, we planned two orthogonal comparisons. The local mismatch effect was investigated through the comparison of the two repeated sequences (ssss and sssS) against the two locally deviant sequences (sssd and sssD) whereas the global mismatch effect was investigated through the comparison of the sequences respecting the block rule (ssss and sssd) against those violating the rule (sssD and sssS). As it was the last tone that defined the different conditions, we restricted the statistical analyses to the 1200ms following the fourth tone.

#### Hypothesis testing

Given the previous literature, we selected a-priori time windows to analyze the mismatch response [200-400ms] and [800-1000ms] to test the late response. Because not all participants had data in the four experimental cells, we used mixed models (lme4; Bates et al., 2015) in R (R Core Team, 2019), which are recommended to deal with missing values. After the mean MEG data were log-transformed to assure a normal distribution, the data were tested in a different model for each time window with participant ID as a random effect and the local and the global status of the condition as two fixed effects (each one with two levels). An additional model was built to explore the effect of individual conditions with participant ID as random effect and condition as fixed effect with four levels. This model was used to focus on post hoc comparisons only, as the main effect of condition is tightly linked to the effects of local and global status. Model significances were tested with a type II Analysis of Variance with Satterthwaite’s method to estimate effective degrees of freedom (lmerTest; Kuznetsova et al., 2017). The combination of restricted maximum likelihood fitting of the model (default in lme4) and Satterthwaite approximation is following Luke’s recommendations (2017) in order to optimally control for type 1 errors. Normal distribution of residuals was assured with a Quantile-Quantile Diagram. Post hoc testing of the models was done by calculating estimated marginal means. The significance level was set to α=0.05, in case of post hoc testing, the significance level was corrected for multiple comparisons by false discovery rate (fdr).

#### Exploratory analysis

As an addition to the hypothesis-driven analyses, we performed an exploratory analysis on the global effect. We compared the time traces of global standards (mean of ssss and sssd) and global deviants (mean of sssD and sssS) after the onset of the fourth tone with a 10000 fold permutation analysis. For this analysis only the participants with data available in all four stimulus conditions (N=15) were considered. In the permutation analysis, the labels (global standard/global deviant) within each participant were randomly shuffled and the difference between time traces of these randomly generated groups calculated. From all permutations, a distribution of differences for each time point was determined. The positions in the time traces where the real difference between groups lay outside the 95^th^ percentile of this distribution were marked as significant differences.

Two time windows of interest were extracted from the results of the permutation test and we computed post-hoc paired tests between the four conditions by calculating estimated marginal means. Then we explored the influence of inter-individual differences – i.e. the newborns’ behavioral state, measured by HRV as a proxy – in this subset of data. We split the participants into extreme groups by taking the highest third and lowest third of recordings of each block. This kind of split was performed to ensure the dominance of one or the other behavioral state during the recording.

Statistical testing was done with mixed models in R as described in the previous section and post hoc testing was done by calculating estimated marginal means. To avoid a circular analysis, we did not include the main effects of local and global rules or conditions. We further studied the impact of behavioral state on the brain responses in general by comparing the event-related-responses (ERRs) to the first tone of all included sequences between infants with low vs. high HRV. We also compared the power spectral density of brain responses between these two groups. These comparisons were done with a permutation test as described previously with the difference that labels were permuted not only within participants but completely at random. Significance level was set to α=0.05 in all cases. Due to the explorative nature of the analysis, no correction for multiple comparisons was applied.

## 3. Results

### 3.1. Hypothesis driven analyses

In the early time window (200-400ms) – assigned to the MMR – analysis showed a highly significant effect of the local rule violation (F(1,47.96)=12.81, p<0.001; Figure 2). There was no effect of the global rule violation and no interaction between local and global rule violations in this time window. When looking at all individual conditions, there was a significant effect of condition (F(3,47.64)=4.76, p=0.006) whereas post-hoc testing revealed that only the sssD deviant was significantly different from both standards (ssss - sssD: t(46.1)=-3.05, p_fdr_=0.011; sssS - sssD: t(48.6)=-3.19, p_fdr_=0.011). The responses for the local deviant (sssd) were weaker but in the same direction (ssss-sssd: t(48.6)=-1.76, p_fdr_=0.127 and sssS - sssd: t(46.1)=-2.003, p_fdr_=0.102; Figure 2). No significant effect was observed in the late time window (800-1000ms).

**Figure 2:**
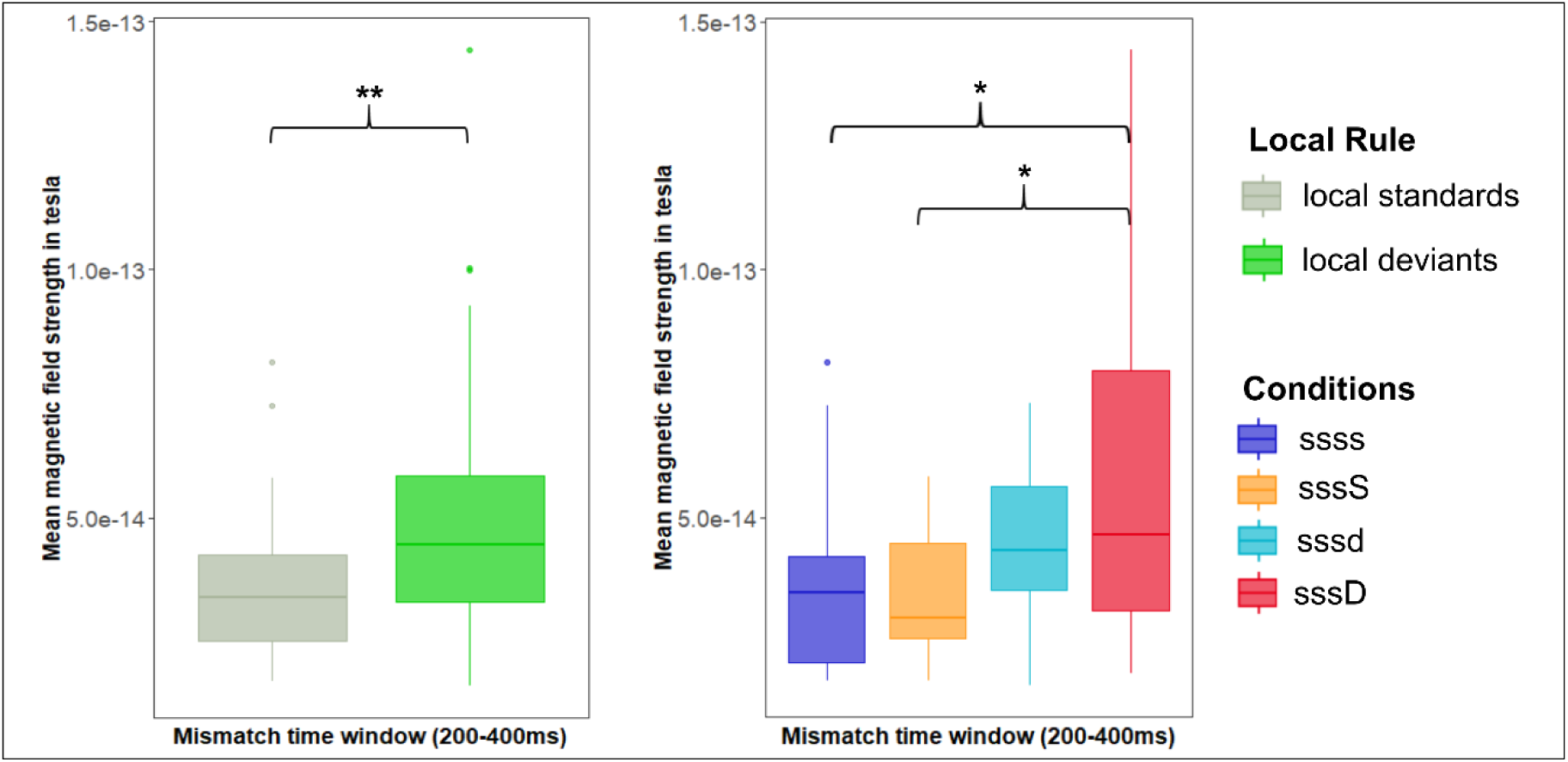
MMR in time window 200-400ms. Left: comparison between local standards and local deviants. Right: individual conditions. * p<0.05, **p<0.001 – fdr corrected. Figures depict original values, while log-transformed data were used for analyses.

### 3.2. Exploratory analysis of the global effect

To further explore the responses to global rule violations, we performed a permutation test on the time series of the 15 infants with data in both types of blocks. This analysis revealed two significant time windows during which conditions with global rule violations (sssS and sssD) differed from global standards (ssss and sssd; Figure 3A). Inter-individual variance within these time windows (80-115ms and 720-745ms) can be seen in Figure 3B. We thus further examined whether the newborns’ behavioral state was affecting the global effect. We considered the HRV, measured by SDNN as a proxy for it.

**Figure 3:**
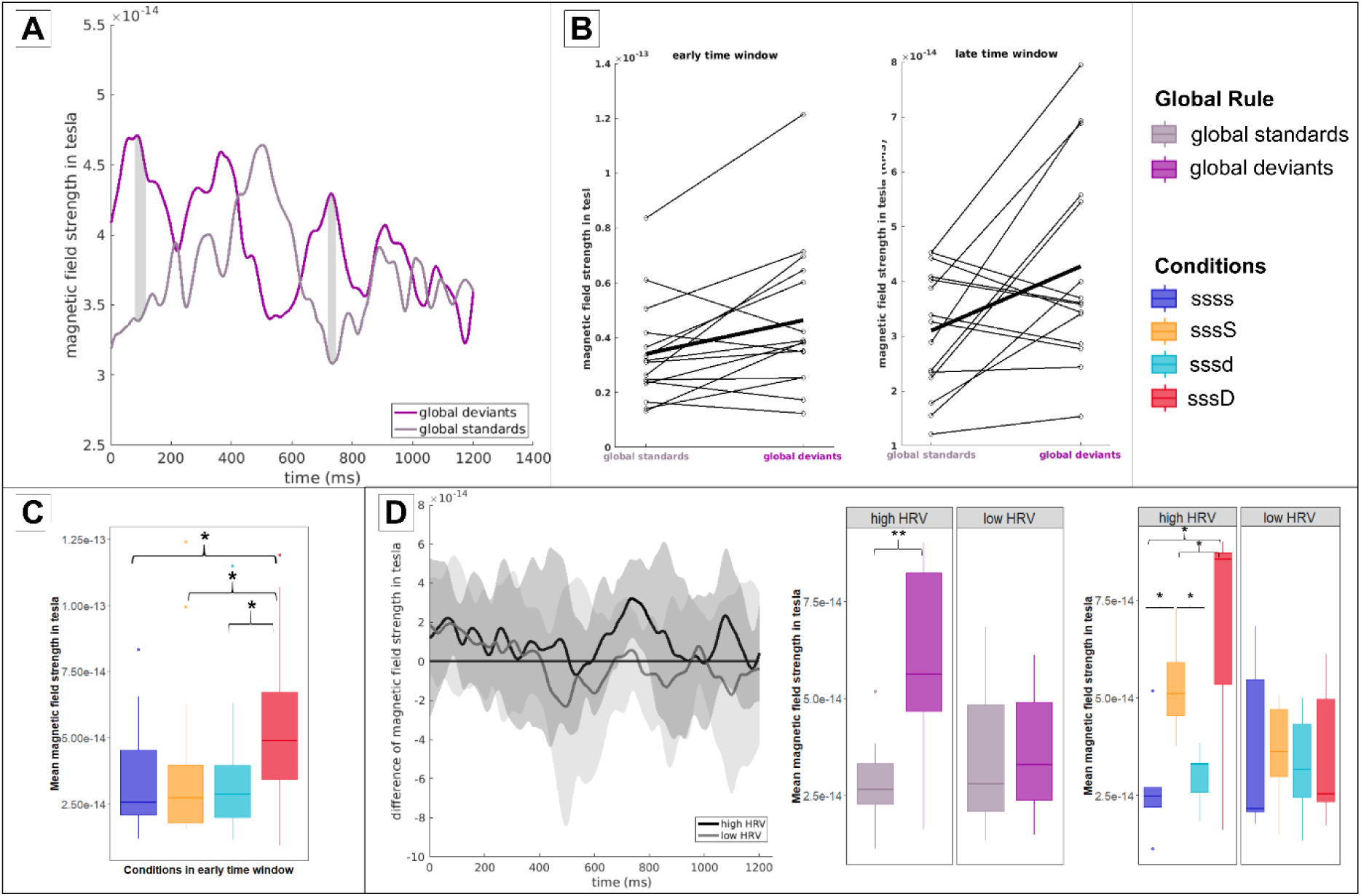
A: results of permutation test. Comparison of global standards and global deviants. Grey areas depict significant differences (p<0.05) B: Data variability in early and late significant time window. Each line represents mean data in that window of one participant, bold line the average. C: Data in early time window (80-115ms) split by condition * depict significant differences (p<0.05). D: Data in late time window (720-745m). Left: differences between global deviants (sssD & sssS) and global standards (ssss & sssd) split by HRV. Shaded area marks standard deviation. Low HRV group contains lowest third of values, high HRV group highest third. ; Middle: Interaction global rule with HRV; Right: Conditions split by HRV groups * depict significant differences (p<0.05); ** p<0.001. Figures depict original values, while log-transformed data were used for analyses.

Our participants displayed a SDNN range from 9.37 to 27.79ms (M=19.01; SD=5.09; Figure 4A). We split them into two extreme groups (N=5 each, M=24.04ms (SD=2.53) in the high HRV group and M=13.4ms (SD= 3.33) in the low HRV group). Those values are in accordance with the SDNN values of active and quiet newborns reported in previous studies (Doyle et al., 2009; Lucchini et al., 2017). Figure 4B shows that those two groups already differ in their brain response towards the first tone of a sequence, which is independent of condition – with a larger ERR in high HRV participants. Additionally, brain signals in the frequency domain of participants with a low HRV showed a significantly lower power spectral density compared to high HRV participants (Figure 4C).

**Figure 4:**
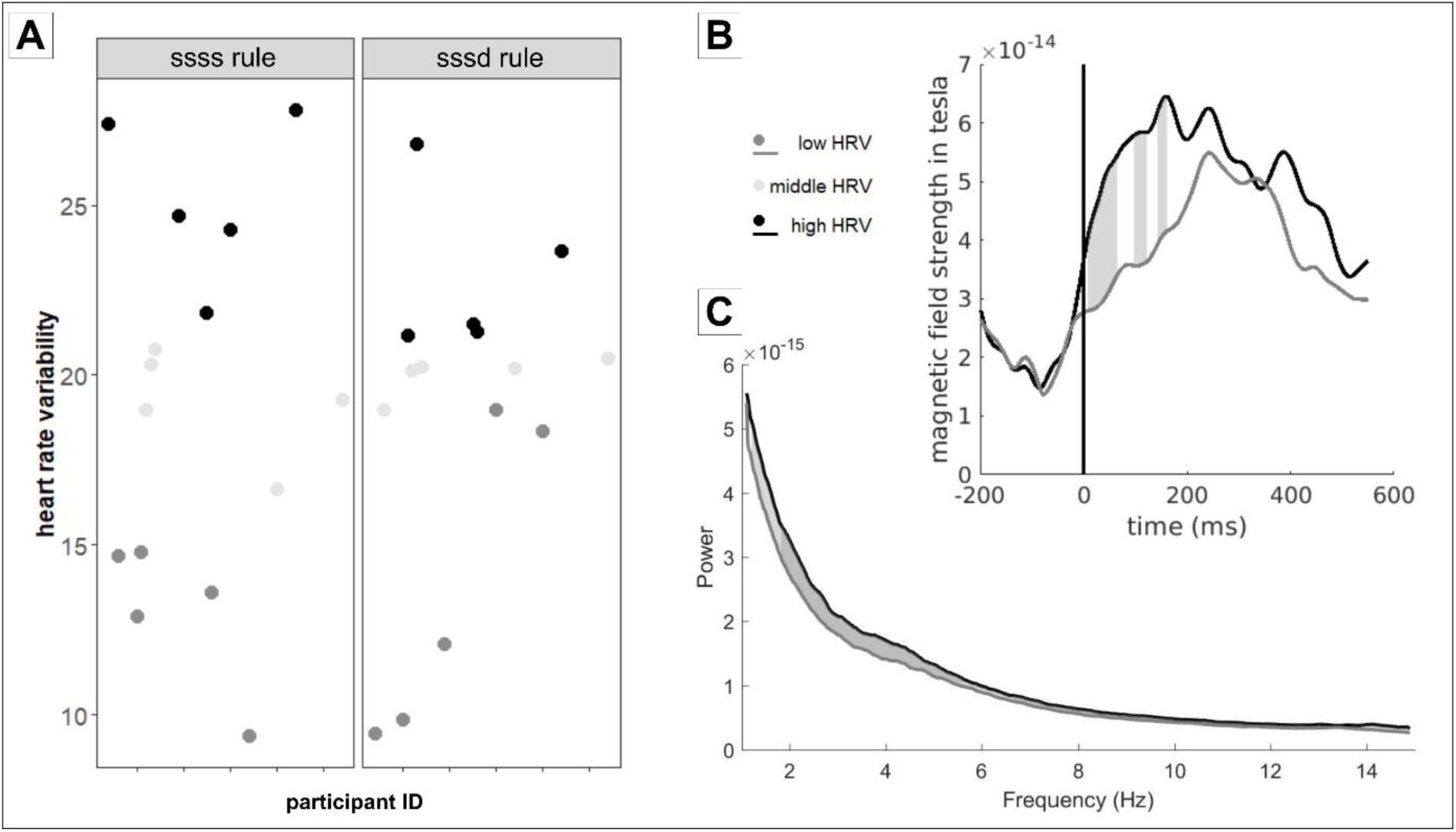
HRV values for n=15 participants included in the exploratory analysis. A: Distribution of values in recording blocks with ssss rule and sssd rule. B: Event-related-response towards first tone of sequence (all conditions included) split by HRV groups. C: Power spectrum of participants in high and low HRV groups over recording. Grey areas depict significant differences (p<0.05), dark grey p<0.01.

We then analyzed the impact of the newborn’s behavioral state on the global effect, there was no effect in the early time window. Post-hoc tests revealed that only the local and global deviant (sssD) differed from all other conditions (sssD - ssss: t(42)=2.74, p=0.009; sssD - sssS: t(42)=2.36, p=0.023; sssD - sssd: t(42)=2.75, p=0.009; Figure 3C). Thus the effect was mainly carried by the large response to the rare deviant tone (rare sssD sequences among many ssss sequences). On the contrary, in the late time window (720-745ms), the split by HRV revealed a significant interaction between HRV and the global rule (F(1,25.54)=11.69, p=0.002; Figure 3D). Newborns with high HRV displayed a large difference between global standards and global deviants (F(1, 10.93)= 41.44, p<0.001) whereas this difference was not significant in low HRV newborns (F(1,9.54)=0.11, p=0.747). Post-hoc tests revealed that each globally deviant tone induced a significantly larger response than the global standard tones in the high HRV condition (sssS - ssss: t(25.4)=3.21, p=0.004; sssD - ssss: t(21.2)=4.34, p<0.001; sssS - sssd: t(21.2)=3.05; p=0.006; sssD - sssd: t(25.4)=3.24,p=0.003). The two local conditions as well as the two global conditions did not differ (sssd - ssss: t(25.4)=0.56, p=0.582; sssD - sssS: t(25.4)=0.59, p=0.563). Conditions in the low HRV group did not differ significantly. Figure 3D illustrates the time-series of the global effect in the low and high HRV states.

Finally, we calculated a Pearson correlation between HRV and the global effect in the late time window in each type of blocks across all 20 participants, not restricting the analysis to those selected into the high and low HRV groups. In case of the block with the ssss rule, there was a significant correlation between the global effect (ssss-sssD) and HRV (cor=0.55, p=0.022). In the block with the sssd rule, the correlation (sssd-sssS with HRV) was in the expected direction but not significant (cor=0.32, p=0.19).

## 4. Discussion

In this study, we presented two hierarchical levels of auditory deviance to newborn infants. A local change within a four tones sequence and a second-order violation relative to the global structure of a recording block.

Our results show a clear MMR towards local rule violations within the time window (200- 400ms) expected in infants (Friederici et al., 2002; Basirat et al., 2014). However in the late time window (800-100ms) when responses towards global rule violations were expected, no significant difference was observed. To further explore responses to a second-order violation, we performed a permutation analysis that isolated two time windows of interest (one early and one late). The early difference was mainly due to the rare deviant tone (rare sssD sequences among many ssss sequences). This sequence induced a larger response than all the other sequences. The late time window (720-745ms) in which both rule-deviant sequences tended to evoke a larger response than both rule-standard sequences fits to results by Dehaene-Lambertz and Dehaene (1994) who reported a late frontal negativity as early as 680ms in 2 to 3 month-old infants. Compared to their findings, the time window of interest found in our study was rather narrow and subsequent analysis showed that this response was affected by newborns’ behavioral state.

### Early responses reflect a local computation

Given the experimental paradigm, a local effect is defined by a violation of tone repetition, so the sssd and sssD sequences should differ from the two ssss and sssS sequences. Mismatch responses following a sound change in a series of repeated stimuli have been robustly found in newborns, and even before term in preterm neonates (Mahmoudzadeh, Wallois, Kongolo, Goudjil, & Dehaene-Lambertz, 2017) and in fetuses (Draganova et al., 2005; Draganova et al., 2007). Responses to sssd, although in the same direction, were weaker than for sssD. This implies that the comparison was not limited to one sequence but that the tones’ occurrence was accumulated over several sequences. In this case, the deviant tone is less frequent in a block with a ssss rule than in a block with a sssd rule. Thus, the deviant tone in a sssD sequences is very unexpected, evoking a large error signal. In 3-month-olds, the ERR amplitude during the MMR time window was also proportional to the occurrence of the two sounds in the block and not limited to the sequence, with the following order: sssD > sssd > sssS > ssss (Basirat et al, 2014). A similar result was observed by King et al. (2013) in the local-global paradigm in adults and Baldeweg et al. (2004) reported, that the amplitude of mismatch responses increased with decreasing probability of a deviant stimulus among standard stimuli. Thus the prediction error is modulated by the probability of occurrence of each tone (Friston, 2005) and MMR results can be put well into the context of the probabilistic updating model proposed by Wacongne et al. (2012).

The large response to the sssD sequence therefore explains the early difference (80-115 ms) captured by the permutation analysis which was exclusively related to this condition, whereas in the 200-400 time window, the local effect was more distributed over the two locally deviant sequences. The early latency of this response compared to the expected latency might be related to the simple auditory features of a pure tone compared to the more complex speech stimuli used in Friederici et al. (2002) and Basirat et al. (2014). Mismatch responses in newborns in a time window that early have previously been found for white noise stimuli (Kushnerenko et al., 2007). The finding that this very early response was not influenced by the behavioral state is congruent with the assumption that this mismatch response is a mainly automatic process.

### Late responses and second-order computations

A global effect is demonstrated when both the rare sssS and sssD sequences elicit a different brain response than the frequent ssss and sssd sequences, although the sssS sequence only contains repeated tones. Both sequences violate the rule of the block constructed by the 30 learning trials and the (75 %) rule congruent trials during the testing phase. The late time window where permutation testing revealed a significantly higher response towards the two rare sequences compared to the frequent sequences was rather narrow and showed a high variance. However, from previous studies we know that the late response towards the global deviant in the local-global paradigm vanishes during sleep (Strauss et al., 2015). Additionally, the LSW response in infants has until now only been shown in awake infants (Friederici et al., 2002). Because in a passive listening paradigm in newborns it is hard to distinguish between sleep and wakefulness, especially since newborns easily switches from one state to the other within a few minutes, we used HRV to account for behavioral states. To ensure actual differentiation between active and quiet newborns, we used the extreme groups (upper third and lower third) of our sample. Of course, the selection of extreme groups within our sample left us with a very low sample size, yet this split ensures a more pronounced difference between the infants’ states. The exploratory analysis showed that the behavioral state has indeed an effect on the processing of second-order regularities. Infants with a higher HRV displayed a clear signature of this process, as both rare sequences were significantly different from the frequent sequences. This suggests that our analysis over the whole group was probably not very sensitive. Figure 3D additionally illustrates, that the global effect was quite long in the high-HRV group. This relation of global effect and HRV did still hold when all 20 infants were considered, especially in the ssss block in which the sssD violation is easier to detect.

Infants with high HRV might have been awake or in active sleep or more rapidly fluctuating between stages. In any case, the fact that the MEG power spectral density was higher and moreover that the amplitude of the ERR to the first tone was larger, supports the hypothesis that they were more in line with the sound stimulation. Already in fetuses, significantly faster ERRs were recorded when they were in an active state compared to a quiet state (Kiefer-Schmidt et al., 2013). Although Strauss et al. (2015) did not detect global rule violations in adults in REM sleep, Raimondo et al. (2017) were able to detect responses towards global rule violations in minimally conscious patients by showing an elongation of the timing of the heart beat following a global rule violation. This shift in timing could not be detected in patients in an unresponsive wakefulness state nor was it shown for local rule violations in any group. Their research emphasizes the connection of the central and autonomous nervous system whereby they speculate that patient’s minimal attention toward the stimuli induced the heart acceleration following the second-order rule violation.

As newborns spend a lot of time in active sleep during their first weeks of life, we may wonder about the role of this state early in life, especially given the amount of learning taking place during this period. First, it might be possible that short periods of arousal are intertwined without overt behavior as it is the case in minimally conscious patients and thus newborns might be less sleeping than thought. These periods would allow infants to analyze their environment and start to learn. Second, the quality of the sleep itself might be different. Wilhelm et al. (2013) showed that children exposed to sequences of lights were better than adults to transfer this implicit encoding into explicit rule knowledge after sleeping. The improved performance was mediated by stronger hippocampal activations in children compared to adults. Similarly, Friedrich et al. (2015) have reported that after sleep, infants (9-16 months of age) were better in generalizing a word’s meaning to new exemplars of the category. Although these particular learning examples were dependent on slow wave sleep and not REM, we do not know enough about the early functions of sleep during the first weeks of life to characterize what happened during our recordings. Moreover, Cheour et al. (2002) showed that newborns were better to discriminate between subtle vowel distinctions if they have been exposed to them during sleep compared to a control group without training. These few results point out the need of better monitoring infants’ behavioral state during neural recordings to be able to explore its relation with learning.

In the current study, a bias towards sleeping participants exists due to the nature of the MEG measurement, as active awake newborns often not complete a full recording block. If they complete a recording block, their data are often excluded during data processing because of a large amount of high amplitude artifacts due to movements. For the measurement of active, awake newborns other techniques like EEG as in Basirat et al. (2014) or functional near infrared spectroscopy (Emberson et al., 2019) could be more suitable as newborns can be measured on their parents’ lap and sensors are attached to the head. On the other hand, MEG has the advantage that newborns can just be placed on the device and no long preparation is necessary.

In summary, we found that human infants, in the first weeks of life, show abilities of hierarchical rule learning, at least when they are in an active state. In the framework of the local-global paradigm, this formation of a memory trace can be interpreted as a basic form of conscious processing as it shows the ability to dynamically adapt to the environment, which is seen as a prerequisite for primary consciousness (Edelman, 2003). These results are in line with those reported by Basirat et al (2016) in 3-month-olds by showing that the MMR is based on sound occurrences spanning more than one sequence, thus implying memory-integrating events over at least a few seconds. However, discovering second-order regularities is a slow process evoking a response after 700ms and needs at least some attention, even if it fluctuates. Therefore it is possible to measure hierarchical rule learning with a passive listening paradigm in infants. However, a close monitoring of participants’ behavior is crucial for a meaningful interpretation of results. More generally, further studies are needed to explore the role of vigilance and sleep in infants’ learning.

## 5. Author Contributions

*Julia Moser:* Investigation, methodology, formal analysis and writing – original draft. *Franziska Schleger:* Conceptualization, methodology and writing – review & editing. *Magdalene Weiss:* Investigation and data curation, project administration. *Katrin Sippel:* Methodology and software. *Ghislaine Dehaene-Lambertz:* Validation and writing – review & editing. *Hubert Preissl:* Supervision and writing – review & editing. All authors read and approved the final version of the manuscript.

### 6. Acknowledgements

We want to thank Janina Einsele for her support with data acquisition. This work was funded by the FET Open Luminous project (H2020 FETOPEN-2014-2015-RIA under agreement No. 686764) as part of the European Union’s Horizon 2020 research and training program 2014– 2018 and the German Federal Ministry of Education and Research (BMBF) to the German Center for Diabetes Research (DZD e.V. 01GI0925). GDL is funded by the European Research Council (ERC) under the European Union’s Horizon 2020 research and innovation program (grant agreement No. 695710).

